# Vascular regional analysis unveils differential responses to anti-angiogenic therapy in pancreatic xenografts through macroscopic photoacoustic imaging

**DOI:** 10.1101/2024.05.27.595784

**Authors:** Allison Sweeney, Marvin Xavierselvan, Andrew Langley, Patrick Solomon, Aayush Arora, Srivalleesha Mallidi

## Abstract

Pancreatic cancer (PC) is a highly lethal malignancy and the third leading cause of cancer deaths in the U.S. Despite major innovations in imaging technologies, there are limited surrogate radiographic indicators to aid in therapy planning and monitoring. Amongst the various imaging techniques Ultrasound-guided photoacoustic imaging (US-PAI) is a promising modality based on endogenous blood (hemoglobin) and blood oxygen saturation (StO_2_) contrast to monitor response to anti-angiogenic therapies. Adaptation of US-PAI to the clinical realm requires macroscopic configurations for adequate depth visualization, illuminating the need for surrogate radiographic markers, including the tumoral microvessel density (MVD). In this work, subcutaneous xenografts with PC cell lines AsPC-1 and MIA-PaCa-2 were used to investigate the effects of receptor tyrosine kinase inhibitor (sunitinib) treatment on MVD and StO_2_. Through histological correlation, we have shown that regions of high and low vascular density (HVD and LVD) can be identified through frequency domain filtering of macroscopic PA images which could not be garnered from purely global analysis. We utilized vascular regional analysis (VRA) of treatment-induced StO_2_ and total hemoglobin (HbT) changes. VRA as a tool to monitor treatment response allowed us to identify potential timepoints of vascular remodeling, highlighting its ability to provide insights into the TME not only for sunitinib treatment but also other anti-angiogenic therapies.

## 1. Introduction

Pancreatic cancer (PC) is the third leading cause of cancer-related deaths in the United States with a median survival time of 4.6 months and 5-year survival rate of 12%[1, 2]. Surgical resection coupled with systemic chemotherapy is currently the only curative treatment option for patients[3]. This poor prognosis can be attributed in part to the asymptomatic nature of PC that leads to late-stage detection, leaving only 10-20% of those diagnosed eligible for surgical resection[4]. Borderline-resectable PC must rely on preoperative therapy to shrink tumors prior to resection to improve surgical outcomes[5, 6] and neoadjuvant chemotherapy has been shown to increase survival, improve the chance of a full resection, and reduce the frequency of positive margins following surgery[4, 7]. Despite this, a considerable proportion of patients (∼80%) with PC experience recurrence after undergoing resection and/or neoadjuvant chemotherapy, ultimately resulting in patient death[8, 9]. The biological factors underpinning the aggressive and treatment-resistant nature of PC can in part be attributed to features of the pancreatic tumor microenvironment (TME)[10]. The pancreatic TME is characterized by an abundance of stromal cells and extensive extracellular matrix, while remaining hypovascularized[10-12], resulting in a severely hypoxic TME which significantly influences tumor metabolism, therapeutic resistance, and angiogenesis[10, 12, 13]. Angiogenesis, the formation of new blood vessels, is a crucial element in the growth and spread of solid tumors. The irregular shape and function of tumor vasculature, may result in decreased blood flow and oxygenation, impeding the delivery of therapeutics to the tumor site[14-16].

Angiogenesis inhibitors are a family of therapeutics that act by suppressing the development of new blood vessels and recent research has shown that antiangiogenic treatment may be tailored to normalize tumor vasculature[17-19]. Vascular normalization employs modest doses with brief treatment durations to ameliorate the structural and functional irregularities of tumor blood vessels and sensitize tumor tissue to conventional therapy[17, 20-22]. Enhancing the uniformity of functional vascular density and improved configuration of arteries can lead to a decrease in regions of hypoxia and acidosis[17, 23-25]. In preclinical models, vascular normalization via anti-angiogenic therapies has been demonstrated to boost tumor blood supply and oxygenation[26-28], reduce metastatic burden[29, 30], and enhance the efficacy of ionizing radiation[31-34], chemotherapy[35-38], and immunotherapy[39-42]. The majority of antiangiogenic drugs target the vascular endothelial growth factor (VEGF) pathway, either by inhibiting VEGF-A with neutralizing antibodies or by blocking the VEGF-receptors (VEGFRs) with tyrosine kinase inhibitors (TKIs)[43-45]. Sunitinib is a multi-targeted TKI which utilizes the latter approach, inhibiting the activity of a number of tyrosine kinases, such as VEGFRs, platelet-derived growth factor receptors (PDGFRs), and stem cell factor receptors (KIT)[46, 47]. Sunitinib is authorized for the treatment of a variety of cancers, including renal cell carcinoma and gastrointestinal stromal tumors (GIST) such as PC[48, 49]. Much promise has been shown in administering sunitinib in combination with traditional treatments for PC as sunitinib has been demonstrated to make PCs more sensitive to radiation treatment *in vitro* and *in vivo*[50, 51]. Studies on combinatorial treatment have shown that the co-administration of sunitinib with gemcitabine in orthotopic PC models and nab-paclitaxel in subcutaneous PC models enhanced survival and reduced tumor burden compared to monotherapy[52, 53].

Although rapid innovations in cancer therapeutics have allowed for more targeted destruction of solid malignancies evaluating therapy response remains an obstacle, as there are limited endogenous radiographic indicators to aid in therapy planning and monitoring. Monitoring vascular structure and function is of particular interest, as these factors play a pivotal role in understanding therapy-induced changes in the TME[10, 12, 46]. Research into the functional indicators of therapy response in PC is critical for giving timely, accurate feedback on treatment efficacy and developing strategies for personalized medicine. Ultrasound-guided photoacoustic (PA) imaging (US-PAI) has received a lot of attention in recent years due to its non-invasive capacity to provide spatially co-registered anatomical, functional, and molecular data of the TME using endogenous contrast. PAI combines optical excitation and acoustic detection to generate high resolution images containing functional information about biological tissues[54, 55]. PAI involves delivering nanosecond pulsed light into tissue, which is absorbed by chromophores and converted into heat causing thermoelastic expansion and contraction of the absorber and generation of acoustic waves[56, 57] detectable by US transducers. The minimal scattering of acoustic waves in biological tissues allows this hybrid modality to reap the benefit of increased penetration depth compared to purely optical techniques[54, 58]. Based on wavelength selection, US-PAI can display detailed functional and molecular information for a wide range of endogenous chromophores[59, 60]. The absorption spectra of hemoglobin changes when bound to oxygen, allowing the use of multi-wavelength PAI to assess blood oxygen saturation (StO_2_) and hemoglobin concentration (HbT) by independently measuring oxyhemoglobin (HbO_2_) and deoxyhemoglobin (Hb) distributions[61]. Among approaches used to image vasculature, PAI stands out for its exceptional scalability, making it suitable for imaging in the micro-to macroscopic scales. PAI has shown to be a promising modality to evaluate and monitor response to anti-angiogenic therapies based on changes in vascular morphology and StO_2_, which are strongly associated with tumor hypoxia, according to preclinical investigations in murine models[62-65].

In addition to StO_2_, microvessel density (MVD) has shown a correlation with aggressiveness in a variety of malignancies[66-69]. The emergence of tailored anti-angiogenic medication provides the possibility to use MVD analysis as both a prognostic and therapeutic marker. MVD can be measured with a variety of histological and *in vivo* imaging techniques, including PAI[69-72]. To our knowledge, the use of PAI to measure MVD *in vivo* has been mostly limited to PA microscopy (acoustic and optical resolution) or mesoscopy[72-80]. Microscopy techniques may attain far greater spatial resolution, but they are usually restricted in terms of depths up to 1 mm[81]. In PAI specifically, mesoscopy refers to depths from 1-5 mm, with a resolution in the range of a few to tens of microns[80, 81]. In the push towards clinical translation of PAI, there comes the hurdle of balancing system resolution and penetration depth. Clinical imaging of tumors, specifically volumetric imaging, will require macroscopic configurations for adequate depth visualization. PA macroscopy encompasses depths exceeding 5 mm and offers resolution ranging from tens to hundreds of microns, in which individual microvessels cannot be resolved[82]. This illuminates the need for a surrogate marker to classify relative MVD within a tumor using macroscopic PAI. We chose to investigate the feasibility of a surrogate marker for vascular density in PC xenografts treated with sunitinib. Through histological correlation, we have successfully shown that areas of low and high vascular density (LVD and HVD) can be identified through frequency domain filtering of macroscopic PA images. We utilized this classification and performed vascular regional analysis (VRA) of treatment induced StO_2_ and HbT changes. Our results indicate that sunitinib preferentially reduced StO_2_ and HbT in LVD regions during early treatment timepoints, followed by improved reoxygenation of HVD regions in AsPC-1 tumors. MIA PaCa-2 tumors displayed contradictory results, and VRA did not provide sufficient evidence of vascular normalization, highlighting key differences in the characteristics of the two cell lines. VRA as a tool to monitor treatment response allowed us to identify potential timepoints of vascular remodeling and which tumor areas are more susceptible to pruning or reoxygenation from antiangiogenic therapy, highlighting the ability of VRA to provide key insights into the TME which could not be garnered from purely global analysis.

## 2. Materials and Methods

### 2.1 Cell lines and animal models

All animal studies in this work were approved by Tufts University’s Institutional Animal Care and Use Committee (IACUC). Male homozygous Foxn1^nu^ nude mice (The Jackson Laboratory) were subcutaneously injected with 5 million AsPC-1 or MIA PaCa-2 cells in 100 µL of Matrigel (50 µL of Matrigel + 50 µL of phosphate buffered saline (PBS)) using a 28-gauge insulin syringe. AsPC-1 and MIA PaCa-2 cells were obtained from the American Type Culture Collection and cultured in RPMI-1640 (Roswell Park Memorial Institute) and DMEM (Dulbecco’s Modified Eagle Medium) media supplemented with 10% fetal bovine serum and 1% penicillin-streptomycin (100 U/ml), respectively. All cells were grown in T-75 flask and maintained in a humidified incubator at 37 °C and 5% CO_2_. AsPC-1 and MIA PaCa-2 cells were passaged 1-2 and 2-3 times each week, respectively.

### 2.2 Sunitinib treatment

The sunitinib solution was prepared at a concentration of 20 mg/mL by dissolving 200 mg of sunitinib L-malate (Sigma Aldrich) in 1 mL of dimethyl sulfoxide and 9 mL of corn oil. The solution was repeatedly vortexed and placed in a 50 °C ultrasonic water bath for intervals of 2 minutes each until no visible lumps or particles were present. The sunitinib solution was stored in 4 °C. Mice were treated with 80 mg/kg per day with sunitinib or vehicle for 21 days via oral gavage. The treatment regimen began once tumors reached a volume of approximately 50-150 mm^3^. Mice were split into the following groups: Sunitinib group (MIA PaCa-2: n = 7; AsPC-1: n = 6) and no treatment (NT) group receiving 1x PBS via oral gavage (MIA PaCa-2: n = 8 ; AsPC-1: n = 6).

### 2.3 Longitudinal photoacoustic imaging

The Vevo LAZR-X (Fujifilm, VisualSonics) was used to capture US and PA images, utilizing the MX250S linear array transducer with a 6 dB bandwidth of 15-30 MHz, central transmit frequency of 21 MHz, axial resolution of 75 µm, and lateral resolution of 165 µm. The transducer was coupled to an integrated 20 Hz tunable laser (680-970 nm) via fiber optic cables. Throughout each of the imaging sessions, gain (22 dB for US, 45 dB for PAI) and persistence (20 averages per frame) remained constant. The Vevo LAZR-X Oxy-Hemo mode (750 and 850 nm laser pulses) was used to generate the PA images. The experiment timeline is displayed in Fig. 1 with image acquisition for each mouse starting once tumor volume reached a minimum of 50 mm^3^. Pre-treatment imaging was performed 24 hours before administration of the first dose (Day -1: D(−1)). Imaging during the treatment period was performed at precisely 24 and 72 hours (Day 1: D(1) and Day 3: D(3)) after the first administered dose and continued thrice weekly (Day 6 and beyond: D(6+)).

**Figure 1.**
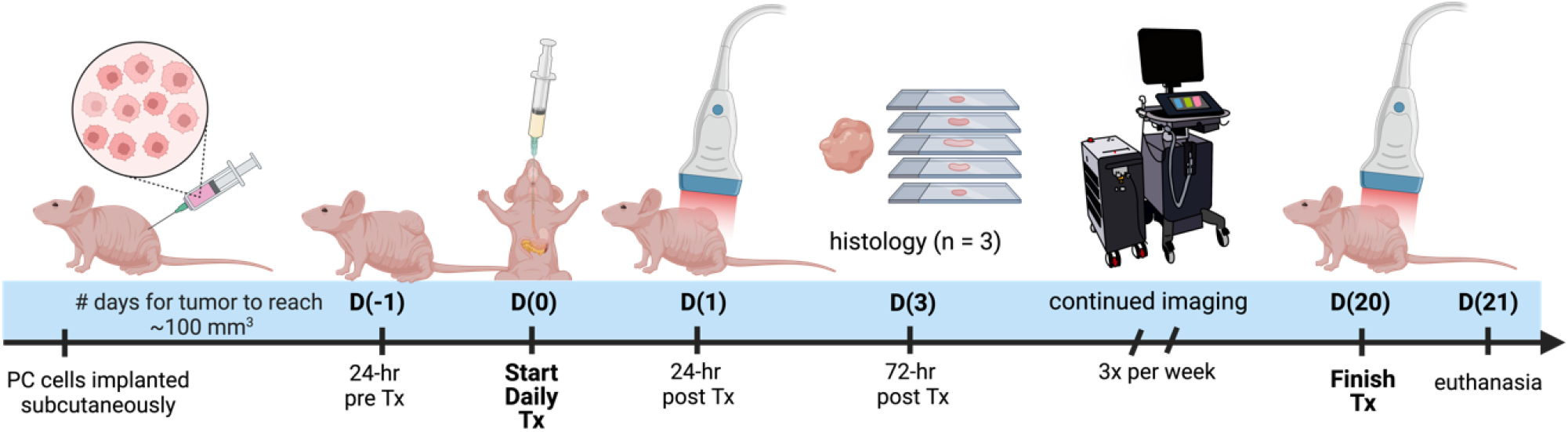
Study timeline describing the imaging, treatment, and histological examination of pancreatic tumor xenografts. D(0) represents Day 0, which is the day of the first treatment or vehicle administration.

The imaging workflow was performed as follows. During the imaging session, mice were sedated with isoflurane (2-3% induction, 1.5% maintenance) with 100% oxygen gas and placed on a heating pad with ECG leads to monitor body temperature, heart rate, and breathing. To improve acoustic transmission between the transducer and the tumor during imaging, a bubble-free ultrasonic transmission gel (Aquasonic100 Ultrasonic Transmission Gel, Parker Laboratories, Inc.) was applied to the tumor. For each frame, an average of 20 images were captured at two wavelengths (750 and 850 nm), yielding a 2D PA image of tumor StO_2_ in roughly 10 seconds. Each dual-wavelength 3D image of a tumor was acquired within 20 to 30 minutes, with the first imaging frame recorded at the back end of the mouse with each succeeding frame moving a 0.15 mm step toward the anterior.

### 2.4 Fluence compensation

Fluence compensation was performed using Monte Carlo simulations utilizing the Monte Carlo eXtreme (MCX) package for MATLAB[83, 84] and the PHotoacoustic ANnotation TOolkit for MATLAB (PHANTOM)[85]. Fluence compensation maps were generated for and applied to all PA images used in analysis. Briefly, 10 million photons (5 million from each fiber) were discharged from the light source toward the tissue volume. For every 0.1 mm^3^ voxel inside the tissue volume, the optical characteristics of absorption coefficient, scattering coefficient, anisotropy factor, and refractive index were assigned dependent upon the tissue type in that area. Different optical properties were assigned for skin, soft tissue, and tumor tissue, which are summarized in Table S1. The localization and labeling of these tissue types within the volume was done utilizing the co-registered US images. In the simulation, water served as the interface between the light source and the tissue, to represent the ultrasound gel used as an acoustic coupling layer between the transducer and tissue. Photon propagation was maintained for 5 ns, which corresponds to the width of a common laser pulse used in PAI. Non-reflective boundary conditions were used throughout the simulation. As the transducer moves throughout the tumor volume to obtain a 3D scan, separate simulations were done for each transducer location and compiled to generate the resulting fluence map. More detailed information on the simulation geometry and parameters used for Monte Carlo parameters can be obtained from Sweeney et al. [85]

### 2.4 Oxygen saturation and hemoglobin imaging via multi-wavelength PAI and spectral unmixing

The wavelength (λ) and depth (z) dependent photoacoustic initial pressure produced by pulsed light stimulation of optical absorbers, assuming stress confinement, may be represented by EQ. 1.

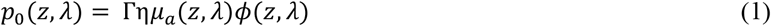

Approximated as constants, Γrepresents the Grüneisen parameter, and ηrepresents the fraction of absorbed light converted into heat. The non-constant *μ*_*a*_(*z*, λ) represents the optical absorption coefficient, and *ϕ*(*z*, *λ*) represents the light fluence. The absorption coefficient is simply the product of the molar extinction coefficient, *ϵ*(*λ*), and concentration, *C*(*z*), of a specific chromophore. In the near infrared (NIR) wavelength range, the absorption of water and lipids is negligible compared to hemoglobin and the mice imaged in this study had insignificant skin pigmentation, allowing us to ignore the contribution of melanin. With the assumption that hemoglobin is the primary biological absorber being imaged, EQ. 1 can be re-written as EQ. 2.

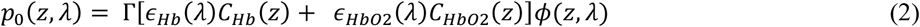

As the two wavelengths used in this study are *λ*_1_ = 750 *nm* and *λ*_2_ = 850 *nm*, The molar concentrations of Hb and HbO_2_, are used to calculate StO_2_ which can be solved for by EQ. 3.

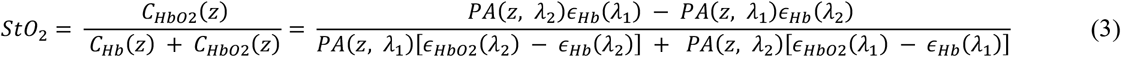

Where *PA*(*z*, *λ*) represents the fluence compensated PA image i.e.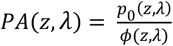.

### 2.5 Image denoising

To improve the signal-to-noise (SNR) of the StO_2_ images, the HbT images were analyzed to find a noise threshold. The noise threshold of each image was calculated using Otsu’s method[86] with two levels. The lower-level threshold was then averaged across all mice for all time-points (Table S2) and rounded to the nearest thousandth to produce a generalized threshold to be applied to all images. If the HbT value of a particular voxel in the full volumetric image was less than the specified threshold, the regions were considered to be avascular, and the corresponding pixel in the oxygen saturation matrix was set to zero and omitted from further analysis.

### 2.6 Vascular Regional Analysis (VRA)

To segment the PA images into regions of HVD and LVD, frequency domain filtering was applied to the volumetric HbT images acquired from spectral unmixing as described in section 2.4 of the text. For clarity, no Fourier analysis was performed on pre-beamformed or single wavelength PA volumes. Volumetric HbT images were normalized to a range of [0,1] and are denoted as I(x,y,z). A 3D Fast Fourier Transform (FFT) was performed on I(x,y,z) to get F(u,v,w) in the Fourier space (EQ. 4) and rearranged by shifting zero-frequency components from the edges to the center of the matrix.

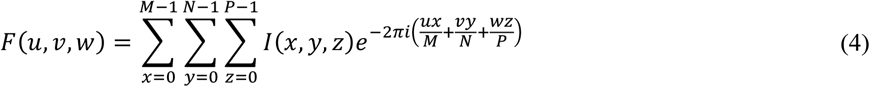

A Gaussian high pass filter (HP) was applied by elementwise multiplication with F to get the filtered volume (F’(u,v,w)) as shown in EQ. 5. The high pass filter was utilized to remove the low frequency components of the image, which we hypothesized would contain signal corresponding to regions of low vascular density.

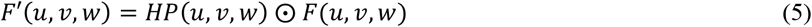

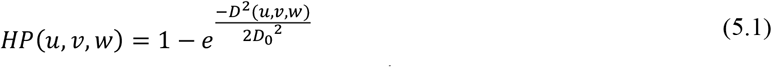

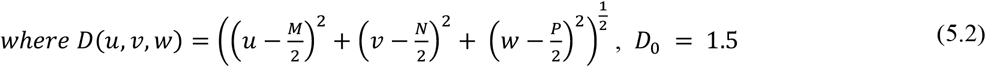

The 2D inverse Fourier transform was used to transform F’ back into the spatial domain (EQ. 6) after rearranging the zero-frequency components back to the edges of the image to get the filtered volume (I’(x,y,z)). The values from the filtered volume, I’, were not used in any data analysis but rather used specifically to segment the relative areas of high vascular density.

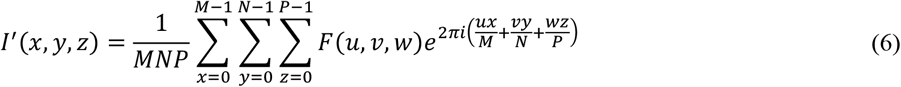

To regionally divide the tumor, the tumor mask was applied to I’, which was then binarized using Otsu’s method[86] to obtain a mask where the foreground represents regions of HVD. For clarity, we are using Otsu thresholding in this context to segment the vascular areas of the tumor and not to omit any regions from analysis. To obtain the complementary mask of LVD regions, HVD was subtracted from the tumor region of interest (ROI). The areas of the tumor considered to be avascular (AV) from the noise thresholding described in section 2.5 are not included in the LVD regions as shown in EQ. 7.

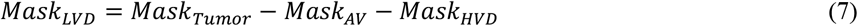

This process allowed the regional analysis of the StO_2_ maps as shown in Fig. 2. As the majority of the low frequency image components were removed with filtering, no analysis was performed on the filtered HbT images. Instead, the filtered images were solely utilized to define the vascular regions, which were then applied to the non-filtered HbT and StO_2_ images.

**Figure 2.**
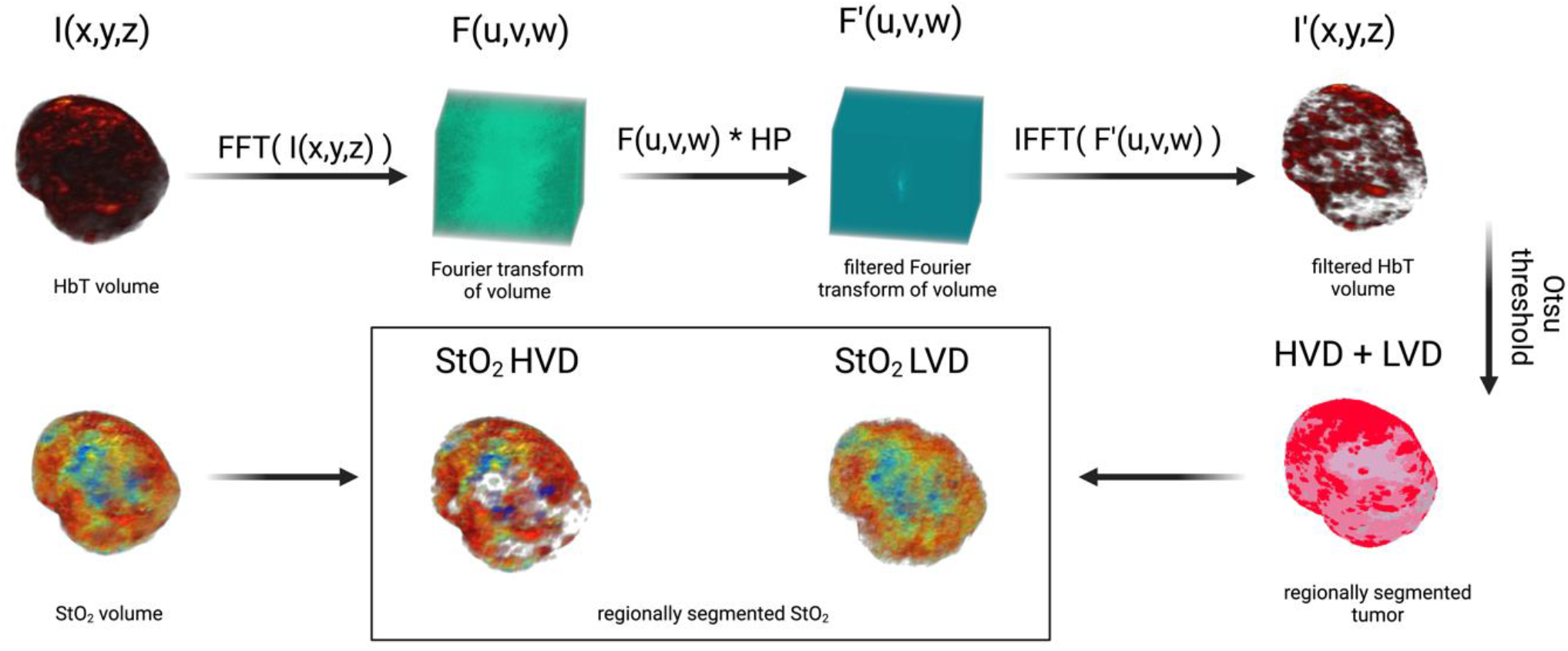
Image processing workflow displaying method for segmenting regions of high vascular density (HVD) and low vascular density (LVD)

### 2.7 Immunohistochemistry

#### 2.7.a Sectioning and staining procedures

Post-euthanasia, tumors were surgically removed with the skin and then placed in optimal cutting temperature (OCT) compound (Tissue-Tek) in the same orientation as the B-scan US-PA images. The tumor tissue was carefully sliced into cryo-sections, each measuring 10 µm in thickness, using a cryotome, then securely affixed to glass microscope slides. The histological examination was conducted using the hematoxylin and eosin (H&E) staining method, as well as immunofluorescence (IF) staining, following a previously reported protocol[63]. Briefly, the cryo-sections were fixed in ice-cold acetone and methanol solution (1:1 v/v) for 10 minutes and then air dried for a duration of 30 minutes, followed by three consecutive 5-minute washes with 1x phosphate buffered saline (PBS). The tissue sections were then blocked with a 1x concentration blocking solution (Blocker™ BSA; #37525, ThermoFisher Scientific™) for 1 hour at room temperature. The immunostaining of vasculature within the tumor sections was performed using a primary antibody, specifically the Mouse CD31/PECAM-1 Affinity Purified Polyclonal Ab (#AF3628, R&D Systems Inc). The antibody was incubated overnight with the tissue sections at a concentration of 10 *μ*g/mL at 4°C. The next day primary antibody was washed off with three rinses of 1x PBS. Secondary antibody Donkey Anti-Goat IgG NL637 Affinity Purified PAb (#NL002, R&D Systems Inc) at a concentration of 20 *μ*g/mL was added to the tissue section and incubated for 2 hours at room temperature. After the incubation, the sections were rinsed in PBS and the nuclei were counterstained and mounted with Slowfade gold antifade mountant containing 4’,6-diamidino-2-phenylindole (DAPI; #S36939, Invitrogen). The slides were imaged at a 4X magnification using EVOS M7000 (ThermoFisher Scientific™) fluorescence imaging system. The IF-stained slides were imaged at the same brightness for CD31 intensity.

#### 2.7.b Immunofluorescence correlation with photoacoustic images

To validate the vascular segmentation algorithm, histological evaluation was performed using the endothelial marker CD31 with a complementary DAPI stain. The corresponding PA cross-section was determined using fiducial markers from the US images and matched to the closest H&E section as previously described[87]. Prior to correlation, IF images were thresholded to a level where autofluorescence was negligible and the tumor region was segmented from the DAPI stain and applied to all IF images. The MATLAB ‘regionprops’ function was used to find the bounding box of the tumor region in the IF images and the complementary bounding box of the tumor region from US-PA images. After cropping all images to the size of their bounding box, the IF images were down sampled to the size of the US-PA images. Thirion’s demons algorithm[88, 89] implemented via the MATLAB ‘imregdemons’ function was used to co-register the down-sampled IF tumor mask with the US-PA tumor mask, and was visually confirmed. Once co-registered, 1 mm x 1 mm rectangular ROIs were drawn, covering the entire tumor region to correlate the average CD31 signal intensity with the fraction of HVD pixels in that same area. Average CD31 signal intensity was calculated as the sum of CD31 intensity in a region divided by the total number of pixels in the region. This parameter was directly correlated with the fraction of HVD pixels, calculated as the sum of pixels labelled as HVD divided by the total number of pixels in the region. For the correlation analysis, 1 cross section was analyzed for 3 different tumors, giving 97 total ROIs, each containing relative amounts of LVD and HVD.

### 2.8 Statistical analysis

GraphPad Prism (La Jolla, CA) was utilized to execute all statistical analyses. Pearson correlation analysis (two-tailed) was performed to validate the vascular segmentation algorithm as described above. The Pearson’s correlation coefficient was calculated for each tumor individually and en masse. Volume growth rates were calculated from exponential fitting of the volume growth curves for each individual mouse. The fit growth rate value between groups were compared using the extra sum-of-squares F test. Growth rate values were omitted from analysis in the case of poor fitting due to non-treatment affects (R^2^ < 0.6). For statistical comparison between two groups at a specific time point, an unpaired two-sample t-test was performed. To statistically compare HVD with LVD parameters, a one-tailed paired t-test was conducted as we were looking to see if there was a significant change in one direction. When comparing more than two unpaired groups (NT and sunitinib treated) and two different cell lines (MIA PaCa-2 and AsPC-1), ordinary two-way ANOVA was utilized (Fisher’s LSD Test). A p-value < 0.05 was considered statistically significant for all analyses.

## 3. Results

### 3.1 Validation of vascular segmentation algorithm with immunohistology

Qualitatively there is a strong visual resemblance between the US image and H&E stain of a representative MIA PaCa-2 tumor as shown by the tumor shape and fiducial markers (Fig. 3A,C, black arrows). The HbT image of the same frame (overlayed onto the US image) and corresponding CD31 stain show excellent visual correlation (Fig. 3B,D). The areas labeled as HVD regions (white arrows) match areas of high CD31 signal, whereas the areas labeled as LVD regions (yellow arrows) match the areas of low CD31 signal intensity. Correlation between the average CD31 amplitude within a region and the fraction of pixels labeled HVD for 3 representative tumors is shown in Fig. 3E. Each point on the plots shown represents a 1 mm x 1 mm ROI. The data points and ROIs that correspond to each tumor are separately plotted in Fig. S1. The Pearson’s correlation coefficient indicates strong correlation between CD31 and HVD each individual mouse (r = 0.925, 0.681, 0.754) and en masse (r = 0.695). P-values for Pearson’s r can be found in Table S3.

**Figure 3.**
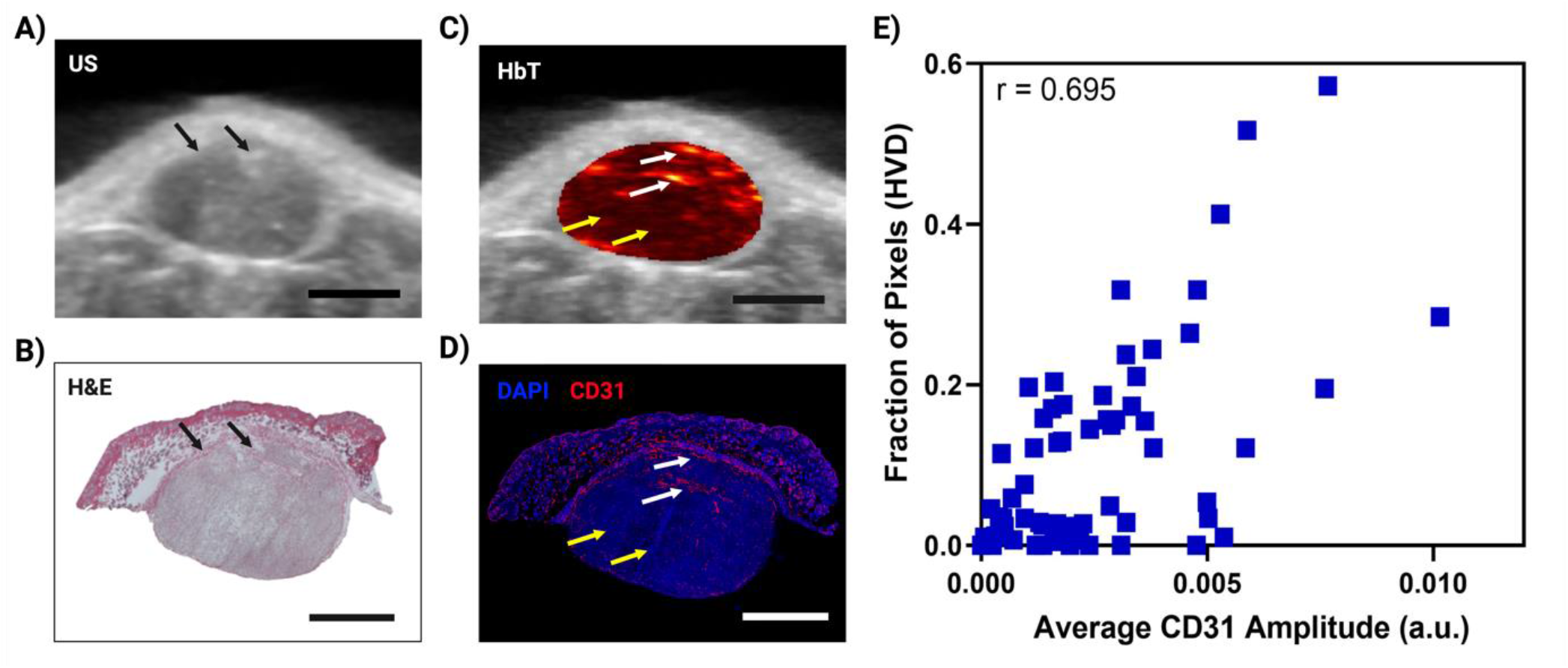
**A)** 2D cross sectional US image in a representative MIA PaCa-2 tumor. **B)** H&E stain of the same tumor cross section shown in A,C. **C)** 2D cross sectional image of HbT overlayed on US in a representative MIA PaCa-2 tumor with white arrows pointing towards areas of HVD and yellow arrows pointing towards areas of LVD. **D)** IF stain of the same tumor cross section shown in A,C with blue representing DAPI and red representing CD31. White arrows pointing towards areas of HVD and yellow arrows pointing towards areas of LVD according to the HbT image. **E)** Plot of average CD31 intensity in a 1 mm x 1 mm ROI versus the fraction of pixels in the ROI labelled as HVD. All scale bars shown represent 2 mm.

To ensure that the segmentation method worked independently of tumor size, vascularity, and treatment regimen, the tumors analyzed all differed in volume (V = 83.9, 105.6, and 246.4, mm^3^), vascular parameters (HbT avg = 0.01, 0.007, 0.006 ; StO_2_ avg = 50.4%, 44.3%, 69.3%,), and encompassed both treatment groups (sunitinib, sunitinib, vehicle). The strong quantitative and qualitative correlation between the pixels labeled HVD and CD31 signal intensity across several tumors’ points to the reliability and repeatability of the proposed vascular segmentation algorithm.

### 3.2 Sunitinib-induced tumor volume changes in AsPC-1 and MIA PaCa-2

Treatment with sunitinib greatly reduced tumor growth rate in both AsPC-1 and MIA PaCa-2 xenografts as shown in Fig. 4A,B. A significant difference in tumor volume between the treated and control groups was apparent within 3 and 8 days of treatment for AsPC-1 and MIA PaCa-2 respectively. Applying an exponential fit to each treatment group reveals that for non-treated tumors, AsPC-1 (k = 0.0635) had a slightly lower best-fit value for growth rate (k) than MIA PaCa-2 (k = 0.0644) despite the fact that non-treated AsPC-1 tumors reached a larger volume by D(20). Our findings were consistent with several studies utilizing subcutaneous pancreatic xenografts, in which quantitative measures of tumor volume showed that AsPC-1 tumors grew larger than MIA PaCa-2[90, 91].

**Figure 4.**
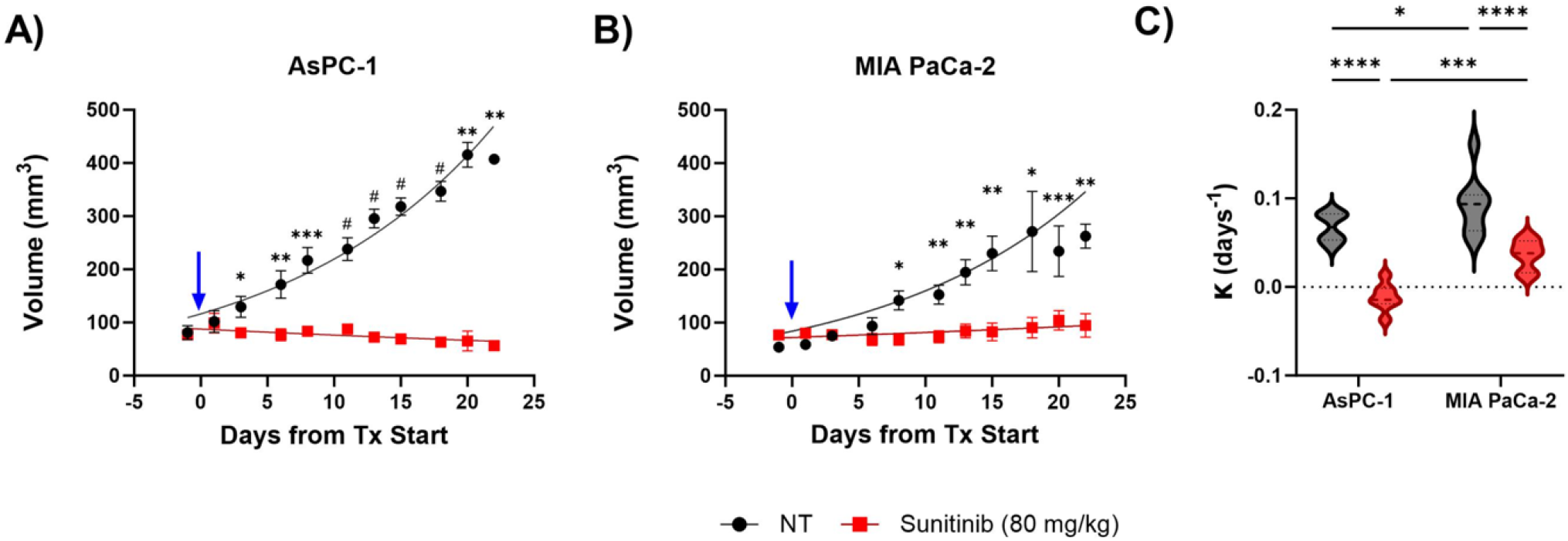
**A-B)** Plot of tumor volume for sunitinib (red) and no treatment - NT (black) groups for AsPC-1 (A) and MIA PaCa-2 (B) tumors with treatment starting at Day 0 (blue arrow). **C-D)** Violin plot of growth rate (C) and volume change from pre-to post-treatment (D) for sunitinib treated (red) and control tumors (black) in AsPC-1 and MIA PaCa-2 tumors. All error bars shown represent SEM. p-values: ^*^ < 0.05, ^**^ < 0.01, ^***^ < 0.001, # or ^****^ < 0.0001

The difference between the average growth rate calculated for each individual tumor is shown in Fig. 4C and reveals that the average growth rates were significantly different between the two cell lines for control (p-value< 0.05) and treated tumors (p-value < 0.001). For both AsPC-1 and MIA PaCa-2, there was a significant difference in the growth rate between sunitinib treated and control tumors (p-value < 0.0001). The observed tumor volume changes throughout the treatment period are shown in Fig. 4D. No significant difference was observed when comparing the two cell-lines for both the control and treated groups. However, within the cell lines, there was a significant difference when comparing the treated and control group for AsPC-1 (p-value < 0.0001) and MIA PaCa-2 (p-value < 0.001). Descriptive statistics reveal that treated AsPC-1 tumor volume reduced by an average of 17.46 mm^3^ (σ = 27.11) and that tumor volume reduction due to sunitinib was consistent between mice. Alternatively, treated MIA PaCa-2 tumors showed an average volume increase of 38.69 mm^3^ (σ = 53.71) during the sunitinib regimen with a much larger variation in response.

### 3.3 Regional vascular response of AsPC-1 to sunitinib

The treatment induced StO_2_ changes in the AsPC-1 cell line are qualitatively and quantitatively displayed in Fig. 5. The respective HbT images for the StO_2_ images shown in Fig. 5 are displayed in Fig. S2. When comparing average StO_2_ of the whole tumor in treated and non-treated groups, a significant difference is seen 24-hours or D(1) (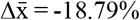, p-value < 0.01) and 72-hours or D(3) post treatment (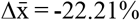, p-value < 0.0001). These findings align with previous work from our group that has shown these timepoints to be significant for StO_2_ reduction in pancreatic xenografts treated with the VEGF inhibitor cabozantinib.[62]

**Figure 5.**
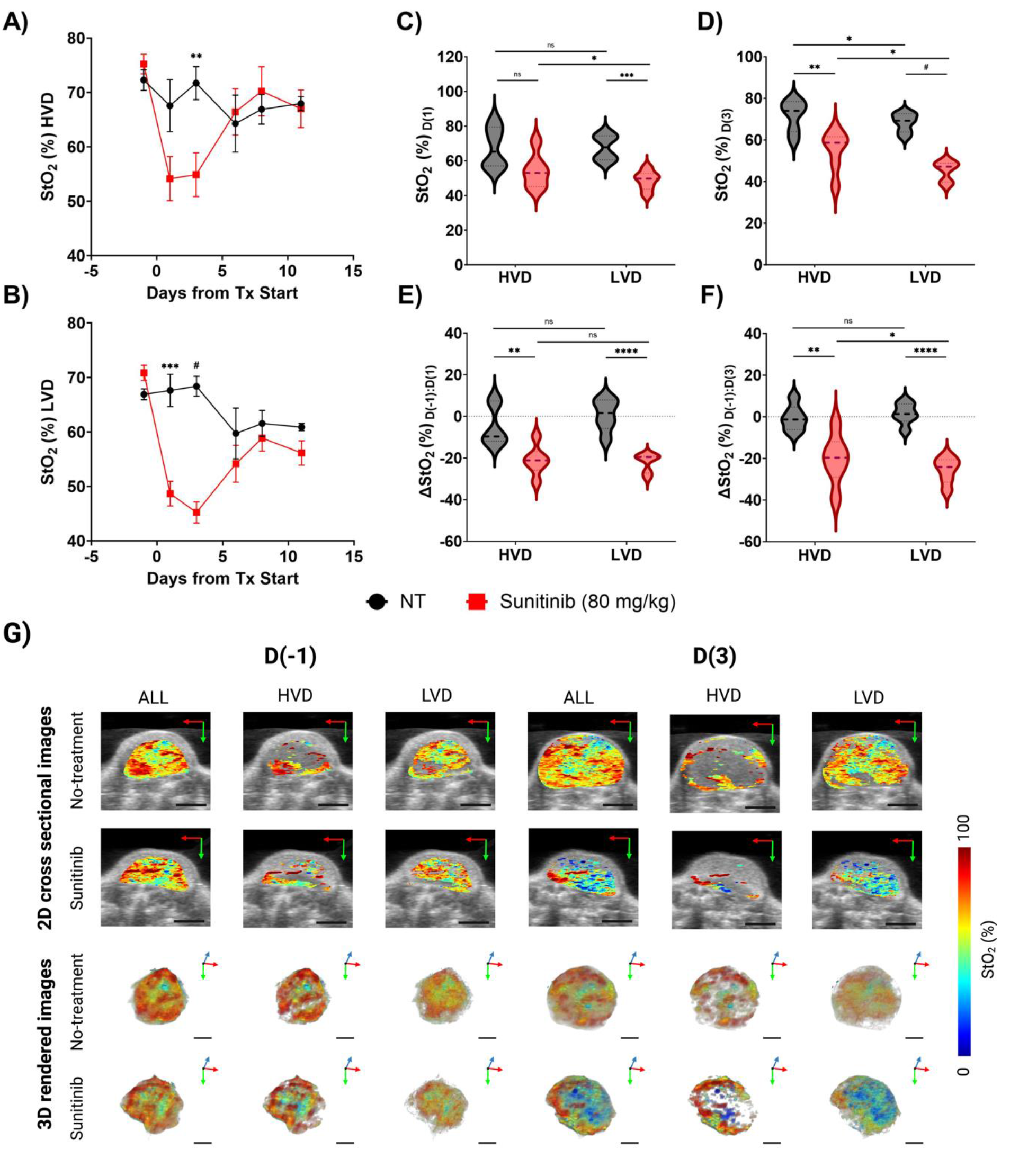
**A)** Plot of StO_2_ in sunitinib treated (red) and no treatment (black) AsPC-1 tumors in areas of high vascular density (HVD). **B)** Plot of StO_2_ in Sunitinib (red) and no treatment (black) AsPC-1 tumors in areas of low vascular density (LVD). **C)** Violin plot comparing StO_2_ on Day 1 (D(1)) for Sunitinib (red) and no treatment (black) AsPC-1 tumors in areas of HVD and LVD. **D)** Violin plot comparing StO_2_ on Day 3 (D(3)) for sunitinib (red) and no treatment (black) AsPC-1 tumors in areas of HVD and LVD. **E)** Violin plot comparing the difference in StO_2_ between Day 1 (D(1)) and Day -1 (D(−1)) for sunitinib (red) and no treatment (black) AsPC-1 tumors in HVD and LVD areas. **F)** Violin plot comparing the difference in StO_2_ between Day 3 (D(3)) and Day -1 (D(−1)) for sunitinib (red) and no treatment (black) AsPC-1 tumors in HVD and LVD areas. **G)** Regional 2D cross sectional images (top) and 3D rendered images (bottom) of StO_2_ in sunitinib (80 mg/kg) and no treatment AsPC-1 tumors displaying the whole tumor (ALL), high vascular density areas (HVD), and low vascular density areas (LVD). All error bars shown represent SEM. p-values: ^*^ < 0.05, ^**^ < 0.01, ^***^ < 0.001, ^****^< 0.0001, # < 0.00001

Segmenting the tumor regions into areas of HVD and LVD reveals that sunitinib is preferentially inducing these StO_2_ changes in LVD areas early in the treatment regimen. As seen in Fig. 5A-C there was no significant difference in StO_2_ between the treated and non-treated tumors at 24-hours post-treatment in HVD areas. Alternatively, there was a strong significant difference between the StO_2_ values of the two treatment groups in LVD areas at the same time-point (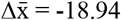, p-value < 0.001). A significant difference in StO_2_ between the HVD and LVD areas was also seen at 24-hours post treatment in the sunitinib-treated group (p-value < 0.05), whereas no significant difference was seen between the vascular regions for the control tumors. By the 72-hour post-treatment time point, significant differences in StO_2_ are present between the treated and control group for HVD (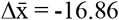, p-value < 0.01) and LVD (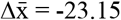, p-value < 0.0001), with the areas of LVD showing a greater difference (Fig. 5D).

Substantial changes in oxygen saturation (ΔStO_2_) from pre-treatment values (Day -1: D(−1)) were seen on D(1) (Fig. 5E) and D(3) (Fig. 5F). The ΔStO_2_ in LVD regions was significantly different between treated to control tumors on D(1) (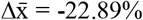, p-value < 0.0001) and D(3) (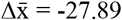, p-value < 0.0001). Significant differences between the treatment groups was also seen for HVD regions on D(1) (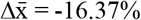, p-value < 0.01) and D(3) (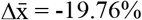, p-value < 0.01), but with a smaller difference between the groups than for LVD regions. There is no significant difference between the LVD and HVD regions for both treatment groups within the first 24-hours of the regimen, however by D(3), LVD areas in treated tumors showed a significantly larger decrease in StO_2_ compared to HVD areas (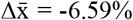, p-value < 0.05), which was not shown in control tumors. Of particular note there was a much larger variation in ΔStO_2_ within HVD areas than LVD areas on D(1) (σ_HVD_ = 12.17%, σ_LVD_ = 7.19%) and D(3) (σ_HVD_ = 7.15%, σ_LVD_ = 4.51%) in the treated mice.

While StO_2_ and ΔStO_2_ are both excellent indicators of vascular changes within the TME[62-64, 92], we also examined the regional changes in HbT to determine if the observed drop in StO_2_ could be attributed to vessel pruning or vascular dysfunction. Regional StO_2_ and HbT values from all treatment days are provided in Fig. S3. The regional HbT changes throughout the treatment period are shown in Fig. S3A-B and shows that sunitinib treated tumors had significantly lower HbT signal at D(1) (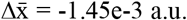, p-value < 0.05) and D(3) (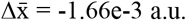, p-value < 0.05) compared to the vehicle group. No significant differences between the treatment groups were seen for HbT in HVD regions at these early timepoints. Given these results, we believe that sunitinib is preferentially targeting LVD areas within the first 72-hours of the treatment regimen in AsPC-1 xenografts.

While the key timepoints associated with treatment-induced StO_2_ and HbT decrease were observed within the first 72-hours of the treatment regimen, the longer-term effect of sunitinib on tumor vasculature was also examined. Specifically, drastic reoxygenation of the HVD regions was observed by D(6) in half of the treated tumors and by D(8) in all but one treated tumor. A paired t-test was performed between HVD and LVD groups for tumors that underwent substantial reoxygenation (ΔStO_2_ > 10%) to determine if the StO_2_ increase was region specific. The amount of reoxygenation that occurred between D(3) and D(8) in these tumors was significant in HVD compared to LVD regions (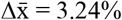, p-value < 0.05). The percent increase in HbT during the D(3-8) time period was significantly different for treated tumors compared to the control in both HVD regions (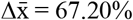, p-value < 0.05) and LVD regions (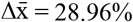, p-value < 0.01). Similar to the regional reoxygenation, HVD regions showed a significantly higher percent increase in HbT than LVD regions (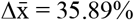, p-value < 0.05) in the sunitinib-treated mice at this juncture.

These observations provide strong evidence that vascular normalization is occurring within the D(4-8) window. From our observations it seems highly likely that sunitinib initially lead to a temporary decrease in tumor StO_2_ while vessels were ablated prior to the onset of vascular remodeling. We see evidence of the preferential ablation of vessels in the LVD regions, indicating these regions housed more immature vessels. Once these immature vessels are pruned, a subsequent increase StO_2_ and HbT is observed, providing surrogate markers for tissue reoxygenation and improved blood flow. These structural and functional changes occur more prominently in the HVD regions indicating the remodeling of more mature vessels in response to the preferential pruning of vessels in the LVD region. Further histological evaluation is needed to support our hypothesis that vascular remodeling is occurring at this period, however all of these observed changes are consistent with previous reports of vascular normalization[17, 23, 24, 93, 94].

### 3.4 Regional vascular response of MIA PaCa-2 to sunitinib

Next, we investigated the MIA PaCa-2 xenografts to determine if the preferential treatment of LVD regions by sunitinib in PC was cell-line dependent. The treatment induced StO_2_ changes in the MIA PaCa-2 cell line are qualitatively and quantitatively displayed in Fig. 6. The respective HbT images for the StO_2_ images shown in Fig. 6E are displayed in Fig. S2. The average StO_2_ of the whole tumor in treated and non-treated groups is significantly different at 24-hours or D(1) (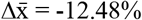, p-value < 0.01) and 72-hours or D(3) post treatment (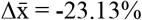, p-value < 0.0001) in the MIA PaCa-2 cell line. Interestingly, performing regional analysis on these tumors reveals the opposite trend as seen in AsPC-1, i.e., in the MIA-PaCa-2 tumors, sunitinib is preferentially inducing StO_2_ changes in HVD areas during early treatment timepoints. There was no significant difference in StO_2_ between the treated and control tumors at 24-hours post-treatment (D1) in LVD areas, while there was a statistical difference in the HVD areas (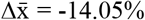, p-value < 0.05), as shown in Fig. 6A-C. Regional StO_2_ and HbT values from all treatment days are provided in Fig. S4. Unlike AsPC-1, there were significant differences between the HVD and LVD areas in the control tumors (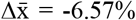, p-value < 0.001), but not in the treated tumors. This indicated differences in angiogenesis between the two cell lines, as at the same timepoint non-treated AsPC-1 tumors showed no differences in oxygenation between these two regions

**Figure 6.**
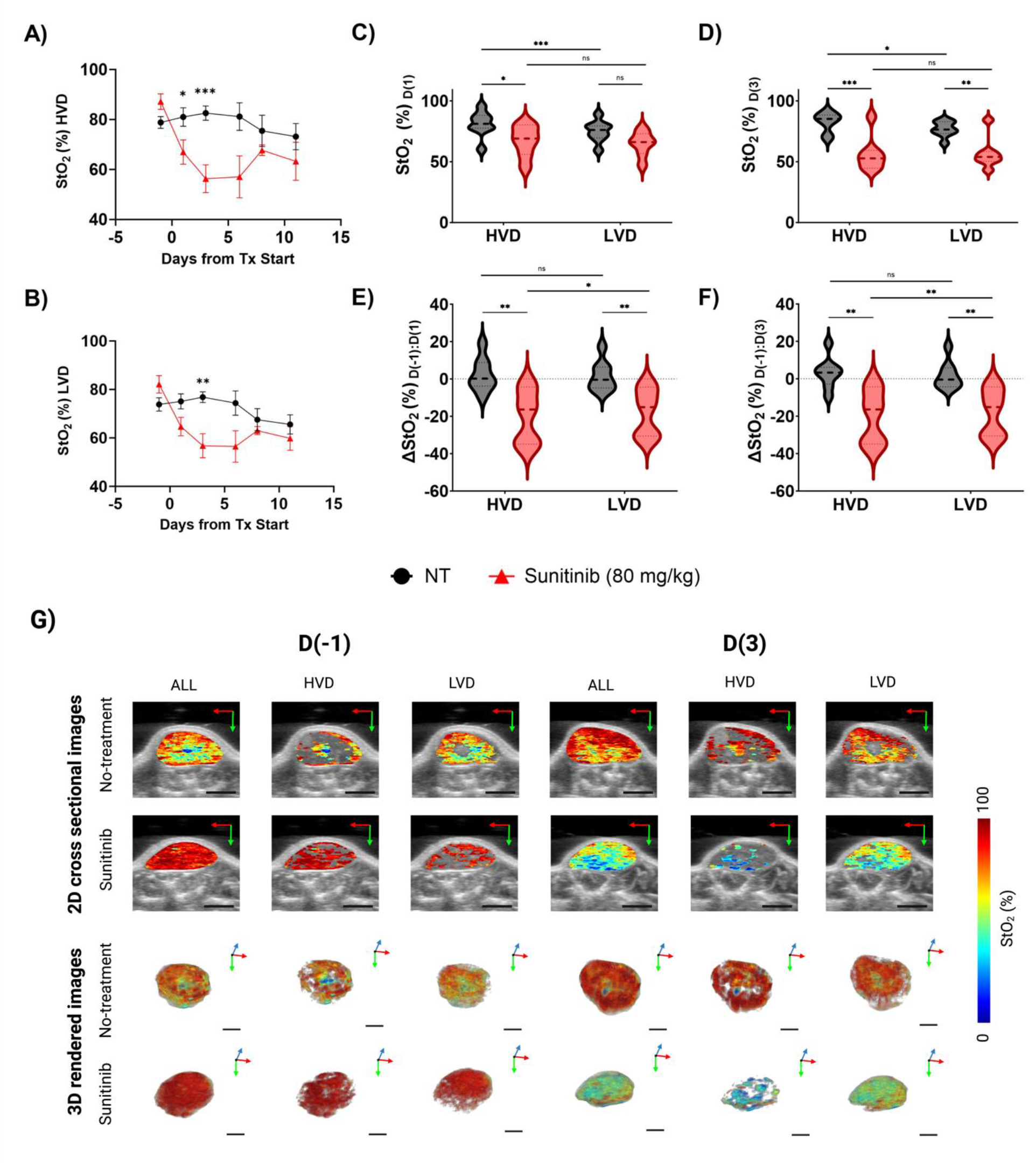
**A)** Plot of StO_2_ in sunitinib (80 mg/kg) treated (red line) and untreated control (no treatment, black line) MIA PaCa-2 tumors in regions of high vascular density (HVD). **B)** Plot of StO_2_ in sunitinib at 80 mg/kg (red line), and no treatment (black line) MIA PaCa-2 tumors in low vascular density (LVD) regions. **C)** Violin plot comparing StO_2_ on Day 3 (D(3)) for Sunitinib at 80 mg/kg (red) and No Treatment (black) MIA PaCa-2 tumors in areas HVD and LVD. **D)** Violin plot comparing the difference in StO_2_ between Day 3 (D(3)) and Day -1 (D(−1)) for sunitinib at 80 mg/kg (red) and no treatment (black) MIA PaCa-2 tumors in areas of HVD and LVD. **E)** Regional 2D cross sectional images (top) and 3D rendered images (bottom) of StO_2_ in sunitinib (80 mg/kg) and no treatment MIA PaCa-2 tumors displaying the whole tumor (ALL), high vascular density areas (HVD), and low vascular density areas (LVD). All error bars shown represent SEM. p-values: ^*^ < 0.05, ^**^ < 0.01, ^***^ < 0.001, #< 0.0001, # # < 0.00001

At the 72-hour post treatment timepoint (D(3)), HbT was significantly lower in the sunitinib group than the vehicle group in LVD areas (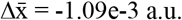, p-value < 0.01), while no difference was shown in HVD regions. StO_2_ between the treated and control groups for both LVD (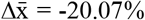, p-value < 0.01) and HVD areas (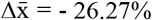, p-value < 0.001) was significantly different as shown in Fig. 6D. When directly comparing the StO_2_ in LVD and HVD areas for each tumor, there were no significant differences between the StO_2_ of these regions in the sunitinib group, but differences were still present in the control group at this timepoint (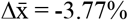, p-value < 0.05) with LVD regions displaying lower StO_2_. As significant differences in StO_2_ between the vascular regions were present in the control group, but not the treated group, we investigated the change in StO_2_ for these regions. From pre-treatment values we investigated ΔStO_2_ for control and sunitinib-treated tumors on D(1) (Fig. 6E) and D(3) (Fig. 6F). The ΔStO_2_ in HVD regions was significantly different between the treatment groups on D(1) (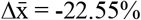, p-value < 0.01) and D(3) (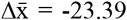, p-value < 0.01). Significant differences between the treatment groups was also seen for LVD regions on D(1) (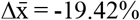, p-value < 0.01) and D(3) (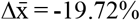, p-value < 0.01), but with a smaller difference between groups than HVD regions.

For the sunitinib group only, ΔStO_2_ was significantly different between the HVD and LVD regions at both D(1) (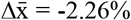, p-value < 0.05) and D(3) (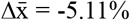, p-value < 0.01). Although the non-treated MIA PaCa-2 tumors displayed differing StO_2_ levels in the HVD and LVD regions, the changes in these regions over time were consistent. Reoxygenation also occurred in the MIA PaCa-2 cell line after the early treatment period, however this reoxygenation was more subtle and was only observed in approximately half of the mice monitored. Additionally, any significant reoxygenation was relatively transient, occurring from D(6-8) rather than D(3-8) as seen in AsPC-1. Similarly to AsPC-1, the increase in StO_2_ was greater in the HVD regions than the LVD regions, however HbT decreased in both the HVD and LVD areas for all treated mice during this window. Neither average HbT nor StO_2_ in sunitinib treated mice was significantly higher than the controls and these values did not increase beyond pre-treatment levels indicating no improvement in vascular function. For these reasons, it is unlikely that any vascular normalization occurred during this window in the sunitinib-treated MIA PaCa-2 tumors. The increase in StO_2_ was relatively small and was only observed for a short time period, leading us to believe it is more likely due to cyclic fluctuations in tumor StO_2_. Towards understanding these effects, our future work will involve monitoring the tumors at more frequent time points during the treatment regimen to investigate the prevalence of short term fluctuations in StO_2_. Additionally, performing contrast enhanced US and conducting oxygen gas challenges will allow us to gauge tumor perfusion.

### 3.5 Vascular differences between pancreatic cancer cell lines

While the effect of sunitinib on tumor volume between the AsPC-1 and MIA PaCa-2 cell lines was comparable (Fig. 4), the regional effect on the vasculature of the two cell lines was markedly different. To understand the discrepancy in response seen, we investigated the baseline vascular characteristics of the cell lines from our *in vivo* work, and relative expression of angiogenic factors from previous *in vitro* work[95, 96]. One of the most crucial proangiogenic factors in cancer is VEGF, which has two primary roles mediated by kinase insert domain receptor (KDR) gene: promoting the growth of new blood vessels (angiogenesis) and increasing the permeability of blood vessels (vascular hyperpermeability)[97-99]. The gene effect scores for KDR and KIT are negative in both cell lines and lower in AsPC-1 (KDR: -0.300, KIT: -0.103) than MIA PaCa-2 (KDR: -0.231, KIT: -0.056), indicating higher dependency of VEGFR and KIT for cell growth in AsPC-1[100, 101]. This aligns well with our findings in which sunitinib reduced the growth rate of both cell lines, but only showed volumetric reduction in AsPC-1.

As shown in Fig. 7A there was no difference in the pre-treatment HbT of the two cell lines when looking at the whole tumor volume, however as seen in Fig. 7B-C the distribution of the vasculature was extremely different between the two cell lines with AsPC-1 tumors displaying a significantly higher fraction of LVD areas and AV areas prior to treatment initiation. Additionally, Fig. 7D displays that the blood oxygen saturation was higher in MIA PaCa-2 tumors and showed much more heterogeneity between tumors than AsPC-1. These results indicate that AsPC-1 tumors may promote less angiogenic activity, and contain more immature microvasculature than MIA PaCa-2, which would align with previous work that has shown MIA PaCa-2 to express higher levels of the pro-angiogenic factors COX-2[95] and VEGF[96] when compared directly with AsPC-1. The expression of VEGF has been shown to positively correlate with high MVD and is a marker for both liver metastasis and poor survival in PC[102]. The higher expression of VEGF in MIA PaCa-2 could be another piece of the puzzle to explain our finding that MIA PaCa-2 tumors display a higher fraction of HVD areas and poorer volumetric response compared to AsPC-1.

**Figure 7.**
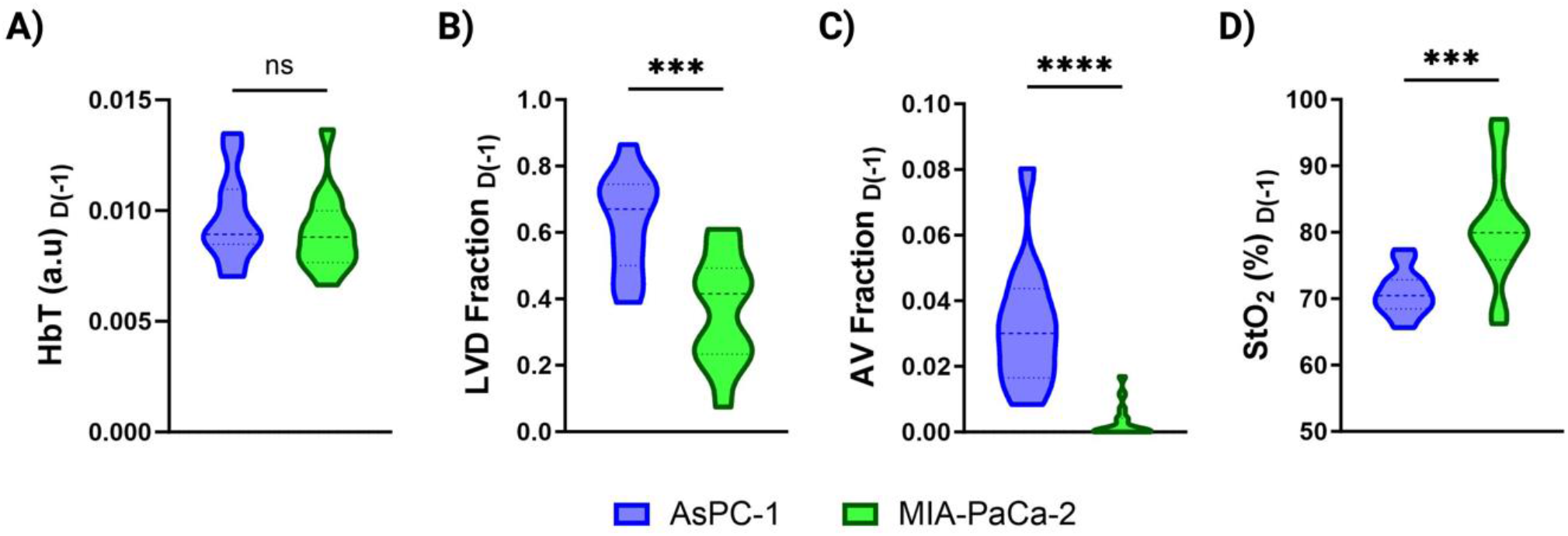
Violin plots comparing the pre-treatment HbT **(A)**, LVD fraction **(B)**, AV fraction **(C)**, and StO_2_ **(D)** in AsPC-1 (blue) and MIA PaCa-2 (green) tumors. p-values: ^*^ < 0.05, ^**^ < 0.01, ^***^ < 0.001, ^****^ < 0.0001

## 4. Discussion and Conclusion

Comprehensive knowledge of the microvascular alterations within a tumor in response to anti-angiogenic therapy is crucial for optimizing efficacy and predicting long-term effects. Several methodologies are currently employed for microvessel segmentation in endogenous contrast PAI such as the threshold-based segmentation [76, 103], morphology-based segmentation[72, 73, 78, 104, 105], and deep learning-based segmentation [75, 79, 106, 107]. While these methodologies are able to directly measure MVD based on microvessel segmentation, they have only been applied to microscopic or mesoscopic imaging configurations [72-80, 103-108] that provide high resolution but lack sufficient penetration depth to enable 3D visualization in a clinical setting. This reiterates the need to develop vascular segmentation or classification methodologies which can be applied to macroscopic, relatively low resolution photoacoustic images. To this end, we successfully developed a methodology for VRA, which is specifically applicable to *in vivo* macroscopic 3D PA images. The VRA methodology is able to differentiate LVD and HVD areas within a tumor to serve as surrogate markers for relative MVD. The VRA methodology was validated with quantitative histological analysis which revealed excellent positive correlation between regions labeled HVD and endothelial marker CD31 (r = 0.695). Once the reliability of VRA was confirmed, we investigated the feasibility of utilizing VRA for monitoring therapy response in solid tumors.

The segmentation methodology was utilized to perform regional analysis on murine pancreatic xenografts treated with the TKI sunitinib. Our findings show that sunitinib induced significant changes in the oxygenation of both AsPC-1 and MIA PaCa-2 xenografts within the first 72-hours of the treatment regimen in agreement with previous literature investigating the effects of anti-angiogenic therapy on subcutaneous pancreatic xenografts[62]. Region-based analysis indicated that sunitinib is preferentially reduced StO_2_ in LVD regions in AsPC-1 tumors and HVD regions in MIA PaCa-2 tumors. At 24-hours post treatment StO_2_ was significantly different between treated and control tumors in only HVD regions for MIA PaCa-2 and only LVD for AsPC-1. At 72-hours post treatment we observed significant differences in ΔStO_2_ when comparing HVD and LVD regions within treated tumors that were not present in untreated tumors. This indicates that the discrepancies seen are due to preferential targeting by sunitinib rather non-treatment effects.

The pre-treatment vascular characteristics of the different tumor models hint towards the superior volumetric response of AsPC-1 to sunitinib compared to MIA PaCa-2. The observation that both AV fraction and LVD fraction were substantially lower in MIA PaCa-2, combined with previous *in vitro* work showing higher pro-angiogenic factor expression in MIA PaCa-2 implies a mature microvasculature compared to AsPC-1[95, 96]. Selective ablation of immature blood vessels from antiangiogenic therapy is a known phenomenon[67, 109] and the preferential targeting of sunitinib could be due to the relative maturity of the microvessels in these regions. Investigation into the mechanisms behind the favorable targeting of specific vascular regions by sunitinib requires in depth analysis that is beyond the scope of this work, but an important future direction. Quantitative assays that measure the level of vascular maturation might potentially indicate the vulnerability of the tumor’s existing blood vessels to sunitinib and would be a logical next step in this work. One weakness of this study is that a subcutaneous model was used, which may not accurately reflect the biology of human PC as much as a genetically engineered mouse model, or orthotopic implantation[110, 111]. This work could also be strengthened by performing an oxygen challenge as done by Tomaszewski et al[92]. An oxygen gas challenge conducted before treatment administration and during the observed reoxygenation window would provide further insight into whether vascular normalization is occurring and the relationship between perfusion and MVD in pancreatic tumors treated with vascular targeted therapies and such methods will be pursued in our future work.

Anti-angiogenic therapies significantly impact several areas of vascular biology and may induce differing levels of endothelial cell death. The impact of anti-angiogenic therapies on tumor StO_2_ is a subject of ongoing debate. Several studies have shown sunitinib therapy to promote vascular normalization by reorganizing the growth of new blood vessels in tumors via removal of dysfunctional vessels[32, 50, 52, 112]. This vascular remodeling can promote the reoxygenation of tumors, showing promise for combination with treatments such as radiation or photodynamic therapy, which require the presence of oxygen[113, 114]. Conversely, other studies have found that sunitinib treatment does not restore normal blood vessel structure and instead leads to hypoxia[115, 116]. Anti-angiogenic therapies which induce hypoxia may render tumors less responsive to the majority of conventional therapies, adding an additional obstacle to dose optimization and timing. The present research did not include any experiments that combined sunitinib treatment with radiation therapy or chemotherapy. Nevertheless, the administration of sunitinib to AsPC-1 xenografts resulted in transient deoxygenation and reduction in vessel density that occurred preferentially within LVD regions. This was followed by reoxygenation in all regions and an increase in blood volume in HVD regions, indicating that the remaining blood vessels had improved functionality within the reoxygenation window. On the other hand, these same metrics showed no conclusive evidence of vascular normalization in MIA PaCa-2 tumors. The differences between HVD and LVD areas highlight the importance of accounting for relative vascular density in measurements of tumor StO_2_ and HbT, and the potential of VRA to provide additional prognostic markers of treatment response and to identify crucial timepoints in which angiogenesis inhibitors can work synergistically with traditional therapeutics.

Overall, our study demonstrates the feasibility of using VRA in macroscopic US-PAI to monitor the vascular microenvironmental changes caused by TKI therapy. VRA coupled with macroscopic US-PAI has the potential to provide valuable insights for the evaluation of key timepoints in which anti-angiogenic therapy is promoting vascular normalization, and how the intratumoral vascular density effects this progression. We have demonstrated the strength and reliability of VRA with endogenous PAI in PC models, however the technology has immense potential to study angiogenesis, monitor therapy response, and inform therapeutic strategy not only in PC but also in several other solid tumors.

## Abbreviations

σ: standard deviation
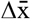: difference between means of two groups
AV: avascular
CD31: cluster of differentiation 31
D(#): day #
DAPI: 4’,6-diamidino-2-phenylindole
DMEM: Dulbecco’s modified eagle medium
FFT: fast Fourier transform
GIST: gastrointestinal stromal tumor
Hb: deoxyhemoglobin
HbO_2_: oxyhemoglobin
HbT: total hemoglobin content
HVD: high vascular density
IF: immunofluorescence
IFP: interstitial fluid pressure
KIT: kinase insert domain receptor
LVD: low vascular density
MCX: Monte Carlo extreme
MVD: microvessel density
NIR: near infrared
NT: no treatment
PA: photoacoustic(s)
PAI: photoacoustic imaging
PBS: phosphate buffered saline
PC: pancreatic cancer
PDGF(R): platelet-derived growth factor (receptor)
PHANTOM: photoacoustic annotation toolkit for MATLAB
ROI: region of interest
RPMI-1640: Roswell Park Memorial Institute
SEM: standard error of the mean
SNR: signal-to-noise ratio
StO_2_: blood oxygen saturation
TKI: tyrosine kinase inhibitor
TME: tumor microenvironment
US: ultrasound
US-PAI: ultrasound-guided photoacoustic imaging
VEGF(R): vascular endothelial growth factor (receptor)
VRA: vascular regional analysis

## Acknowledgements

The authors would like to acknowledge the members of the integrated Biofunctional Imaging and Therapeutics (iBIT) laboratory, specifically Dr. Christopher D. Nguyen, Deeksha Sankepalle, Skye A. Edwards, Ronak T. Shethia, Brooke Bednarke and Avijit Paul and for their useful discussions and support. Funding support from National Institutes of Health grants (S10OD026844, R21CA263694 and R01CA266701) is gratefully acknowledged.

## Contributions

The authors confirm contribution to the paper as follows: conceptualization: S. Mallidi and A. Sweeney; investigation: A. Sweeney, A. Langley, P. Solomon, A. Arora, M. Xavierselvan; formal analysis: A. Sweeney; data curation and visualization: A. Sweeney and S. Mallidi; draft manuscript preparation: A. Sweeney and S. Mallidi; critical revision of the article: A. Sweeney, M. Xavierselvan, and S. Mallidi; project administration and supervision: S. Mallidi; funding acquisition: S. Mallidi. All authors reviewed the results and approved the final version of the manuscript.

