## Supplemental Data for "Vascular regional analysis unveils differential responses to anti-angiogenic therapy in pancreatic xenografts through macroscopic photoacoustic imaging"

| Tissue | $\mu_a$ (mm$^{-1}$)<br>750 nm \| 850 nm | $\mu_s$ (mm$^{-1}$)<br>750 nm \| 850 nm | Anisotropy | Refractive Index |
|---|---|---|---|---|
| Skin | 0.2504 \| 0.2935 | 25.22 \| 21.90 | 0.9 | 1.37 |
| Tumor | 0.0051 \| 0.0061 | 5.278 \| 4.578 | 0.9 | 1.37 |
| Standard Tissue | 0.0110 \| 0.0199 | 10.97 \| 9.418 | 0.9 | 1.37 |

**Table S1**. Optical properties used to fluence compensate PA scans of subcutaneous tumors.

| Day | Mean Threshold | Standard Deviation | N |
|---|---|---|---|
| -1 | 0.00223115 | 4.89E-04 | 28 |
| 1 | 0.00241041 | 4.71E-04 | 28 |
| 3 | 0.00231264 | 4.33E-04 | 27 |
| 6 | 0.00219594 | 3.28E-04 | 20 |
| 8 | 0.0020711 | 4.59E-04 | 19 |
| 11 | 0.00234818 | 0.00056493 | 20 |
| 13 | 0.00226554 | 4.86E-04 | 19 |
| 15 | 0.00223271 | 0.00059685 | 18 |
| 18 | 0.00224655 | 0.00053409 | 17 |
| 20 | 0.00217563 | 0.00056261 | 10 |

**Table S2**. Mean threshold calculated for N mice over all timepoints

| Tumor | HVD vs CD31 amplitude |
|---|---|
| 1 | 5.323E-09 |
| 2 | 6.573E-05 |
| 3 | 3.919E-10 |
| en mass | 3.203e-012 |

**Table S3.** P-values for Pearson's r used in correlation analysis between CD31 and fraction of HVD. Tumor numbers correspond to those shown in Fig S1.

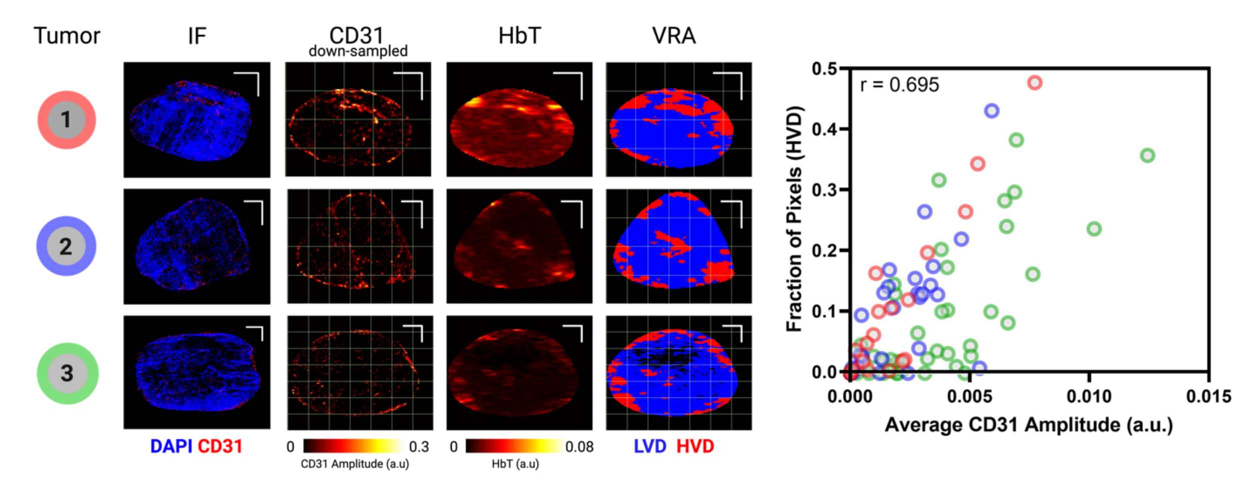


**Figure S1.** Individual tumor cross-sections used to validate VRA algorithm. First image column shows the pre-processed immunofluorescence (IF) image with CD31 displayed in red and the DAPI counterstain displayed in blue. Second column displays the CD31 stain after down-sampling, overlayed with the 1 mm x 1 mm ROIs used for correlation analysis. The third image column shows the matching HbT cross-section input into the vascular regional analysis (VRA) algorithm. The output of the VRA algorithm is shown in the right-most image column and shows areas labeled as high vascular density (HVD) in red, low vascular density (LVD) in blue, and avascular (AV) regions in black. The VRA image is overlayed with the same ROIs used for correlation analysis as shown in the second column of images. All image scale bars represent 1 mm in the x- and y-direction. The average CD31 amplitude is plotted against the fraction of pixels labeled HVD for each 1 mm x 1 mm ROI shown in the down-sampled CD31 images and the VRA images for each tumor.

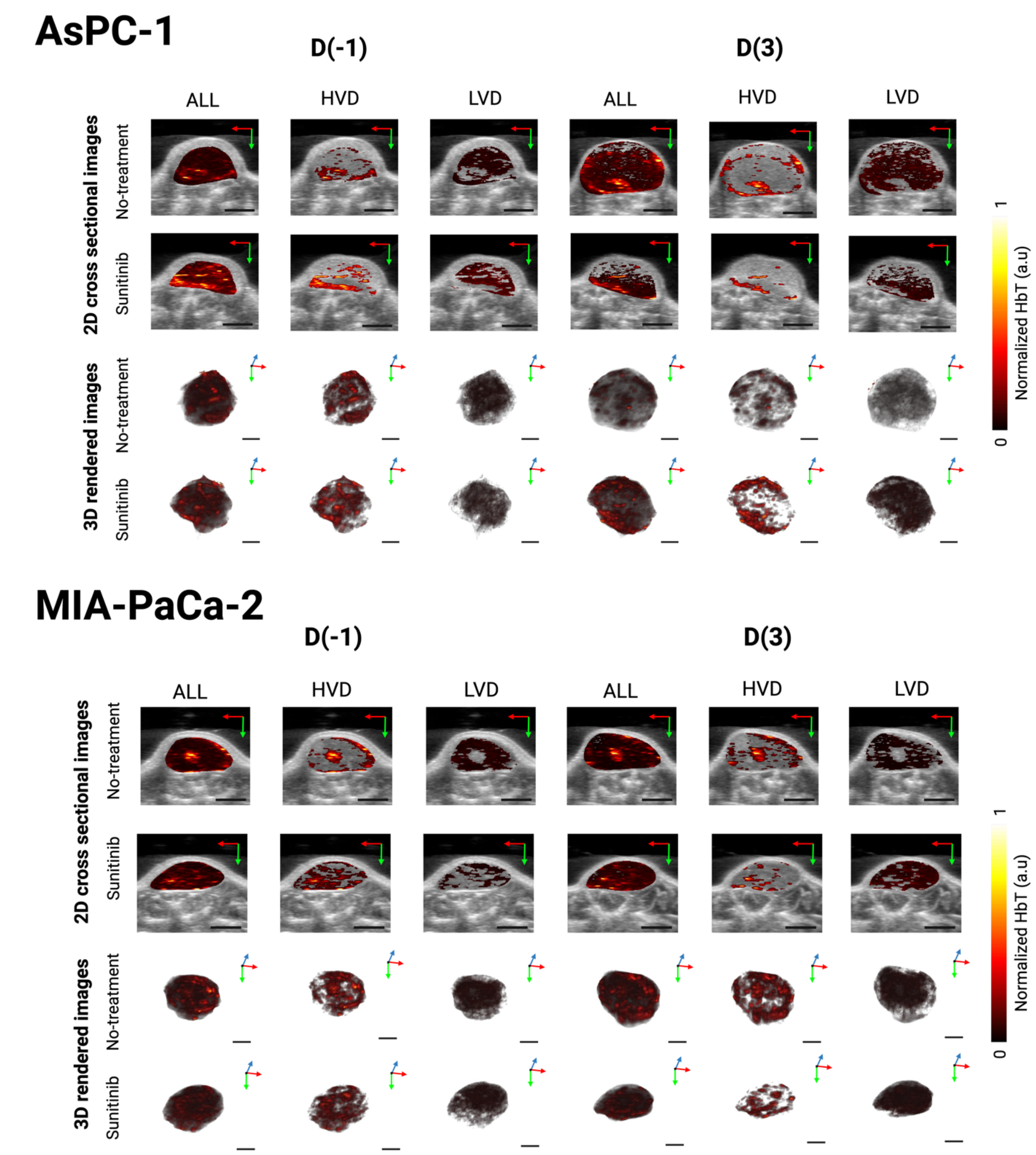


**Figure S2.** Regional 2D cross sectional images and 3D rendered images of HbT in Sunitinib (80 mg/kg) and No Treatment tumors displaying the whole tumor (ALL), high vascular density areas (HVD), and low vascular density areas (LVD). These HbT images correspond to the $StO_2$ images shown for AsPC-1 and MIA-PaCa-2 in Fig 5 and 6 respectively.

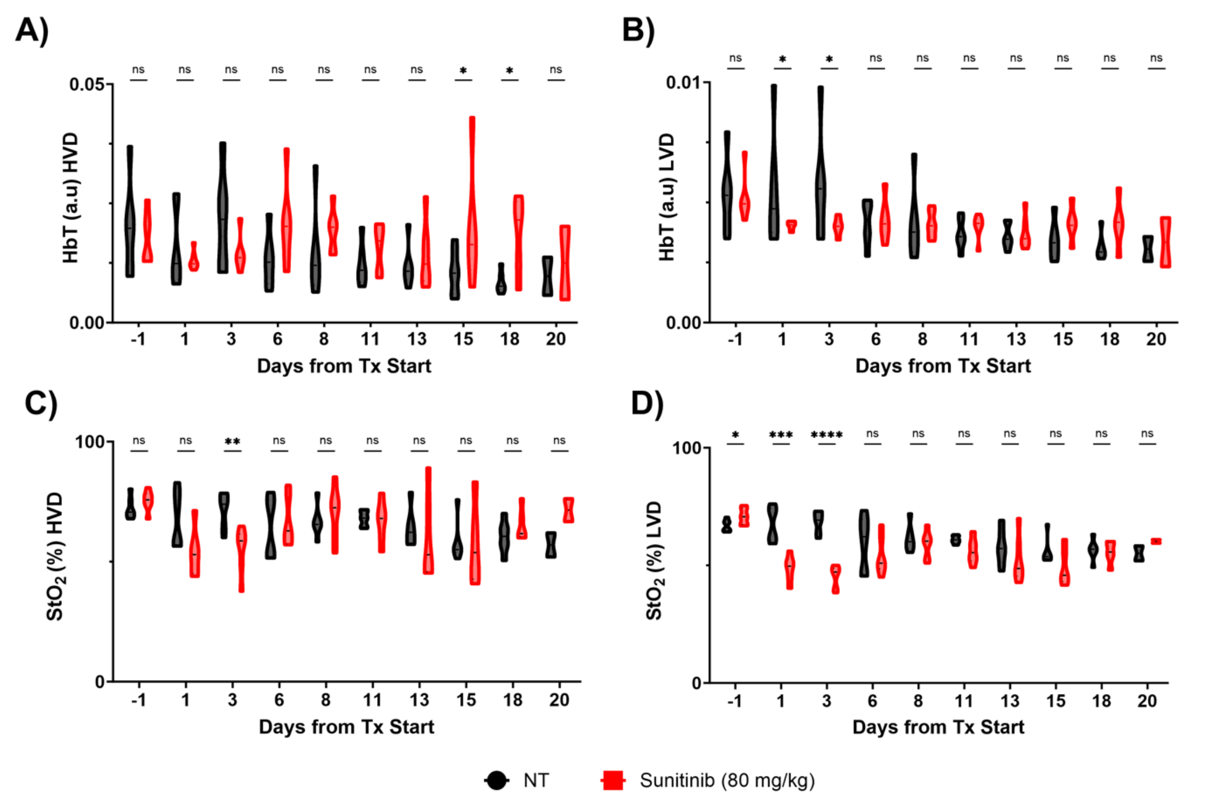


**Figure S3**. **A-B)** Violin plots of HbT in sunitinib-treated (red) and no treatment (NT, black) AsPC-1 tumors in areas of high vascular density (HVD) **(A)** and low vascular density (LVD) **(B)** for D(-1) to D(20)
**C-D)** Violin plots of $StO_2$ in sunitinib-treated (red) and no treatment (black) AsPC-1 tumors in areas of HVD **(C)** and LVD **(D)** for D(-1) to D(20).p-values: * < 0.05, ** < 0.01, *** < 0.001, **** < 0.0001

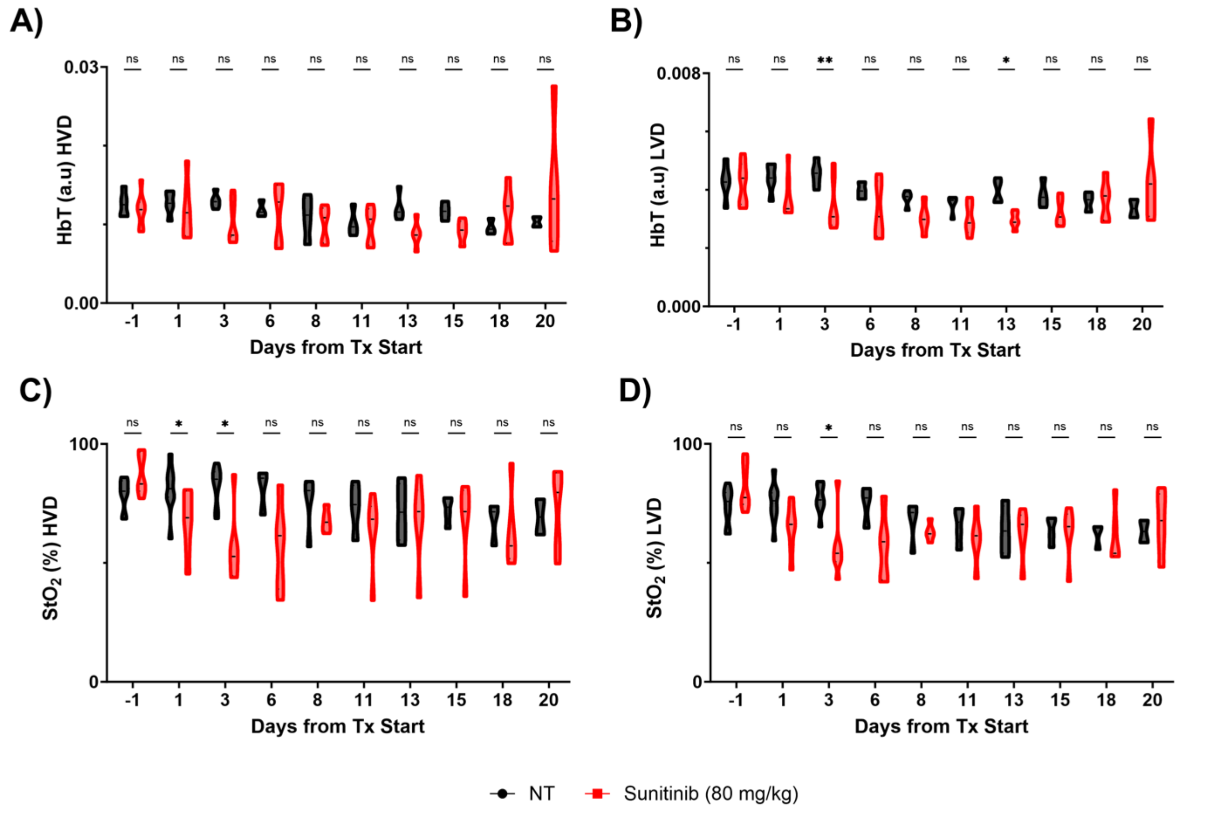


**Figure S4. A-B)** Violin plots of HbT in sunitinib-treated (red) and no treatment (NT, black) MIA PaCa-2 tumors in areas of high vascular density (HVD) **(A)** and low vascular density (LVD) **(B)** for D(-1) to D(20)
**C-D)** Violin plots of $StO_2$ in sunitinib-treated (red) and no treatment (black) MIA PaCa-2 tumors in areas of HVD **(C)** and LVD **(D)** for D(-1) to D(20). p-values: * < 0.05, ** < 0.01, *** < 0.001, **** < 0.0001
